# Linking Electrophysiological Metrics to Oxidative Metabolism: Implications for EEG–fMRI Association

**DOI:** 10.1101/2025.11.04.686568

**Authors:** Xiaole Z. Zhong, Hannah Van Lankveld, Alicia Mathew, J. Jean Chen

## Abstract

Resting-state functional magnetic resonance imaging (rs-fMRI) is widely used to study brain function, yet its biophysiological basis remains incompletely understood. Building on our recent work, we investigated how EEG activity and cerebral metabolic rate of oxygen (CMRO_2_) are related to one another, and how they jointly underpin rs-fMRI metrics. Using a multimodal dataset with macrovascular correction applied to all rs-fMRI metrics, we first examined associations between EEG metrics and CMRO_2_, then applied mediation analysis to evaluate how CMRO_2_ mediates EEG–fMRI associations. We found that bandlimited EEG theta and alphafractional power was significantly associated with CMRO_2_. Bandlimited EEG coherence was also associated with CMRO_2_ across all the bands. Bandlimited EEG fractional power and coherence were also significantly associated with cerebral blood flow (CBF) and oxygen extraction fraction (OEF) in a manner that varied by frequency. EEG broadband temporal complexity was positively associated with CMRO₂ and EEG coherence was negatively associated with OEF. Notably, there are pronounced sex differences in these relationships, which suggests that the biophysical underpinnings of rs-fMRI are sex dependent. Moreover, the baseline metabolic and hemodynamic variables did partially mediate EEG–fMRI associations, with CMRO_2_ serving as the primary mediator. However, most of the mediations are partial, highlighting the complex interplay among electrophysiological activity, oxidative metabolism, and hemodynamics. This study advances our understanding of the biophysical basis of rs-fMRI and provides a foundation for developing sex-specific diagnostic and therapeutic strategies for neurological disorders.

## 1 Introduction

Resting-state functional magnetic resonance imaging (rs-fMRI), most commonly based on the blood-oxygenation level-dependent (BOLD) contrast, leverages spontaneous brain activity (Biswal et al., 1995) to investigate brain function in health and disease. Most rs-fMRI studies focus on connectivity analysis, with BOLD signal power used to characterize the temporal variability of brain signals underlying connectivity measures ((Kannurpatti et al., 2011). More recently, non-linear metrics, particularly entropy, have provided novel insights into the relationship between rs-fMRI signal complexity and cognitive performance (Z. Wang, 2021). Despite the growing number of metrics proposed as markers of neuronal function, their clinical application remains limited, partly due to an incomplete understanding of the neuronal and metabolic mechanisms that underlie the rs-fMRI signal (Lee et al., 2013; Tsvetanov et al., 2021).

Since the early development of rs-fMRI, the preferred approach to clarify the interpretability of FC has been linking it to neuronal activity. Electrophysiological methods such as EEG and MEG are considered gold standards for measuring neuronal contributions. While some human studies have found correlations between EEG and fMRI functional connectivity (FC), results have been inconsistent (Nentwich et al., 2020; Z. Wang, 2021; Wirsich et al., 2020, 2021). EEG power fluctuations may also contribute to FC measures, with associations reported across multiple frequency bands and networks (Liu et al., 2014; Neuner et al., 2014; Phadikar et al., 2025; Scheeringa et al., 2012). Oxidative metabolism (CMRO₂) and glucose metabolism (CMR_glu_) are another key marker of neuronal activity due to the energy demands of neural processes. In particular, CMRO_2_ is intricately linked to fMRI contrast, and rs-fMRI measurements have been linked to CMRO₂ both quantitatively (Raichle et al., 2001) and quantitatively (Zhong, Van Lankveld, & Chen, 2025). However, it is unclear what the link between EEG and CMRO_2_ is in the context of both contributing to fMRI. EEG signals arise from synchronized postsynaptic potentials directed toward scalp electrodes (Malmivuo et al., 1997; Sotero & Trujillo-Barreto, 2008; St. Louis et al., 2016), meaning any neuronal activity not producing synchronized postsynaptic currents is undetectable. Indeed, postsynaptic activity accounts for only ∼74% of total ATP expenditure in humans (Attwell & Laughlin, 2001). Conversely, neuronal activity that does not elicit hemodynamic responses or changes in oxidative metabolism may be invisible to BOLD fMRI, as ATP can be produced via glucose metabolism alone with minimal hemodynamic involvement (Cholet et al., 1997; Vaishnavi et al., 2010). Although these studies have advanced our understanding of the neural underpinnings of FC, their interpretability remains limited, as suggested by our recent series of studies (Zhong, Van Lankveld, & Chen, 2025; Zhong, Van Lankveld, Mathew, et al., 2025).

In our recent studies, we linked FC to both EEG FC (Zhong, Van Lankveld, Mathew, et al., 2025) and CMRO₂ measurements (Zhong, Van Lankveld, & Chen, 2025) individually. While we observed some similar association patterns as reported in previous studies, we also noted some substantial mismatches in both EEG–fMRI and CMRO₂–fMRI association. EEG and CMRO_2_ both measure aspects of neuronal activity, but their biophysical underpinnings differ substantially. As we discussed previously, rs-fMRI reflects oxidative metabolism rather than total metabolism, and accordingly, resting-state CMRO₂ and CMRglu maps are distinct (Markello et al., 2022). One potential source of this discrepancy relates to aerobic and anaerobic glycolysis, which refers to glucose utilization exceeding that required for oxidative phosphorylation despite the presence of sufficient oxygen (Vaishnavi et al., 2010). This may explain why rs-fMRI measurements can diverge from CMRO₂ measurements. Furthermore, although the association between EEG and CMRO₂ is often assumed, it has not been systematically tested.

These considerations indicate that neither the EEG–fMRI nor CMRO₂–fMRI association fully explain the rs-fMRI measurements. Post-synaptic activity, as measured by the EEG, is largely fueled by oxidative metabolism (Attwell & Laughlin, 2001). The EEG–fMRI association is mediated by CMRO_2_ as suggested by previous studies (Attwell & Laughlin, 2001). Conversely, more recent studies suggest discrepancies between EEG and rs-fMRI do not only imply non-neuronal contributions, but may also reflect EEG-invisible neuronal activity that nonetheless consumes oxygen (Tu et al., 2024), highlighting the need to test a CMRO_2_ mediation framework to interpret the underpinnings of rs-fMRI. Furthermore, both CMRO_2_-fMRI and EEG-fMRI associations are strongly influenced by sex as a fundamental phenotype (Zhong, Van Lankveld, & Chen, 2025; Zhong, Van Lankveld, Mathew, et al., 2025). Understanding sex effects is therefore critical for interpreting the role of CMRO_2_ in rs-fMRI findings.

Sex differences in electrophysiological activity have been widely studied, both in terms of power (Corsi-Cabrera et al., 1993; Jausovec & Jausovec, 2010) and coherence (Duffy et al., 1996; Jausovec & Jausovec, 2010). Sex effects are also evident in cerebral metabolism and hemodynamics. Estrogen, the primary female sex hormone, significantly modulates multiple aspects of brain metabolism: it enhances glucose uptake by upregulating glucose transporter expression (Rettberg et al., 2014), increases the total cerebral metabolic rate of glucose (CMR_glu_) by regulating glucose channel activity (Andreason et al., 1994; Rettberg et al., 2014), and promotes aerobic glycolysis by upregulating key glycolytic enzymes (Kostanyan & Nazaryan, 1992; Mannella & Brinton, 2006; Zhu et al., 2023; Znamensky et al., 2003). Consequently, females tend to exhibit higher glycolytic activity and potentially a greater reliance on aerobic glycolysis at rest (Goyal et al., 2019; Rettberg et al., 2014). In addition to metabolic effects, estrogen influences cerebral hemodynamics by modulating vascular tone and perfusion (White, 2002). This aligns with consistent reports that females show higher cerebral blood flow (CBF) but lower oxygen extraction fraction (OEF) compared to males (Aanerud et al., 2017; Gur & Gur, 1990). While findings on CMRO₂ differences are mixed, several studies have reported higher CMRO₂ values in females (Aanerud et al., 2017; Lu et al., 2011). Thus, neither the EEG-fMRI nor the CMRO_2_-fMRI links are generalizable across sexes.

In this study, we acquired a multimodal dataset to investigate associations between EEG and CMRO₂. We also investigate the implications of this relationship for understanding fMRI metrics (**Fig. 1**), which were calculated by incorporating our macrovascular correction pipeline to enhance the neuronal specificity. We examine how these relationships differ between sexes. By characterizing these interdependencies, this study aims to elucidate the electrophysiological and metabolic mechanisms underlying rs-fMRI, thereby advancing our understanding of brain function and informing the development of sex-specific diagnostic and therapeutic strategies for neurological disorders.

**Figure 1.**
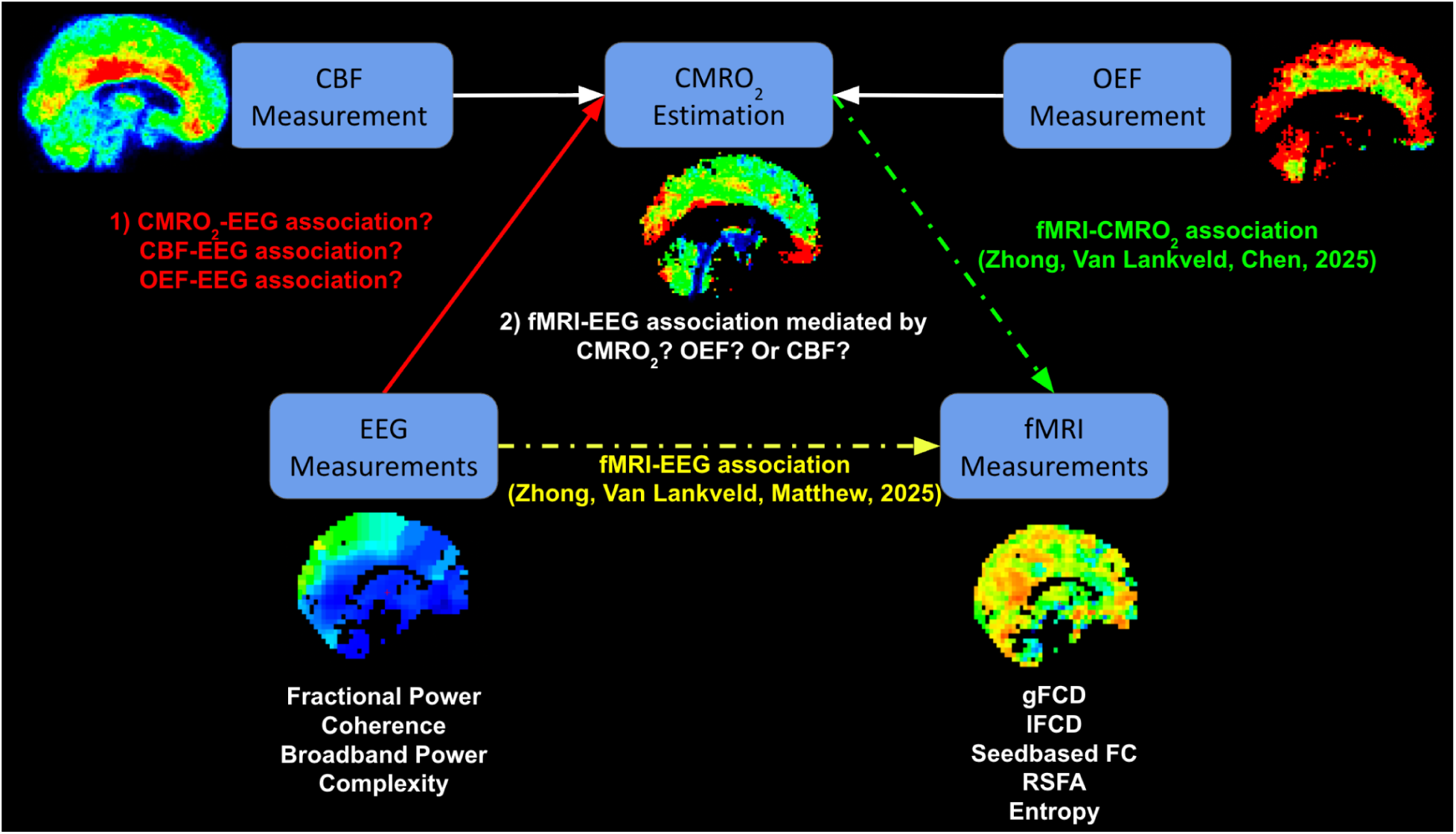
An overview of the analysis framework incorporating the metabolic and electrophysiological associations of rs-fMRI measurements that are tested in this work. In this study, we first examined how EEG measurements are associated with baseline physiological metrics (red arrow). The EEG metrics include bandlimited fractional power, coherence, broadband power, and complexity. The baseline physiological metrics include CMRO_2_ and its correlates CBF, and OEF (white arrows). We further tested here how CMRO_2_, CBF, and OEF mediated (the green arrow) the EEG–fMRI association (yellow arrows). The fMRI metrics include gFCD, lFCD, seedbased FC, RSFA and entropy.

## 2 Method

### 2.1 Participant

A total of 20 healthy participants are involved in this study (10 M/10 F, age = 20 to 32 years). No participant reported having a history of cardiovascular disease, psychiatric illness, neurological disorder, malignant disease, or medication use that may have affected the study. Participants were recruited through the Baycrest Participants Database. The study was approved by the research ethics board (REB) of Baycrest, and the experiments were performed with written consent from each participant according to REB guidelines.

### 2.2 MRI Acquisition

The images were acquired using a Siemens Prisma 3 Tesla System (Siemens, Erlangen, Germany), which employed 20-channel phased-array head coil reception and body coil transmission. The participants were imaged during naturalistic-stimulus viewing to reduce random mind-wandering which increases reproducibility (Gal et al., 2022). The following data were acquired for each participant:

1. T1-weighted structural image (sagittal, 234 slices, 0.7 mm isotropic resolution, TE = 2.9 ms, TR = 2240 ms, TI = 1130 ms, flip angle = 10°).
2. Two time-of-flight (TOF) scans, with coronal and sagittal flow encoding, respectively (coronal: 0.8 x 0.8 x 2.125 mm thickness, 100 slices, TR = 16.6 ms, TE = 5.1 ms, flip angle (α) = 60°; sagittal: 0.8 x 0.8 x 2.125 mm thickness, 80 slices, TR = 16.6 ms, TE = 5.1 ms; flip angle (α) = 60°).
3. One dual-echo (DE) pseudo-continuous arterial spin labelling (pCASL) (courtesy of Danny J. J. Wang, University of Southern California) for recording CBF and BOLD dynamics (TR = 4.5 s, TE1 = 9.8 ms, TE2 = 30 ms, post-labelling delay = 1.5 s, labelling duration = 1.5 s, flip angle (α) = 90°, 3.5 mm isotropic resolution, 35 slices, slices gap = 25%, scanning time = 4 minutes), complete with a M_0_ scan in which the TR was 10 s and the scan time 45 s, all other parameters remaining the same.
4. One multi-echo gradient echo (MEGRE) scan for static R * mapping (TR = 4 s, TE ranged from 5 ms to 60 ms with the step of 5 ms, flip angle = 20°, magnitude and phase recording, same spatial resolution as the pCASL);
5. One SE MRI data for static R_2_ mapping (TR = 10 s, TE =[17, 33, 50, 66, 91, 107, 116] ms, same spatial resolution as the pCASL).

### 2.3 EEG Acquisition

In a separate session from the MRI, a 256-channel Geodesic Sensor Net with a Net Amp 400 amplifier (Magstim Inc., Roseville, MN, USA) was used to record 4 minutes of resting EEG. A naturalistic stimulus video, which is the same as that used during fMRI acquisition, was shown to participants in order to reduce random mind wandering. The EEG electrodes were attached following the international standard 10-20 system, and the impedance of each electrode was controlled per the vendor recommendations (lower than 50 kOhm recommended for the saline-based system). The EEG signal was digitized at a sampling frequency of 1,000 Hz, and the amplitude resolution was set to 0.024 μV.

### 2.4 Macrovascular Correction

The macrovascular correction was performed utilizing a biophysical approach that was developed (Zhong, Polimeni, et al., 2024) and validated (Zhong, Van Lankveld, & Chen, 2025; Zhong, Van Lankveld, Mathew, et al., 2025) in our previous studies. This method was designed to correct macrovascular contributions within and beyond the vasculature, as highlighted in our previous findings (Zhong, Tong, et al., 2024). Since the scan parameters were the same between this study and the previous study (Zhong, Van Lankveld, & Chen, 2025) that applied macrovascular correction, none of the parameters in the correction pipeline had been modified.

A summary of the macrovascular correction pipeline is provided below: 1) generate a vascular mask for each participant and estimate the macrovascular blood-volume fraction (fBV); 2) identify the prototypical venous signal fluctuation from the in-vivo BOLD signal from the voxel with the maximum fBV, which was then demeaned and normalized by its maximum (this signal can be assumed to be driven primarily by T_2_ fluctuations that mimic the variation in venous blood oxygenation, and can also be assumed to contain little information on localized neuronal activity (Zhong, Polimeni, et al., 2024)); 3) simulate macrovascular BOLD signals using our macro-VAN (vascular anatomical network) model; 4) assess the lag between the simulated signal and the in-vivo signal for each voxel (lag that leads to maximum absolute cross-correlation coefficient); 5) regress the simulated signal from the in-vivo signal for each voxel as nuisance regressor (linear regression model).

### 2.5 fMRI Preprocessing and Analysis

Two pipelines were used to process the rs-fMRI data: one for investigating seed-based FC and the other for seed-independent measures. In both pipelines, the first five volumes of BOLD data were rejected to allow the BOLD signal to enter steady state. All rs-fMRI metrics were calculated after macrovascular correction. Moreover, all metrics below were averaged within each functional network listed in the *Seed-based rs-fMRI analysis* subsection to calculate network-wise metrics.

#### 2.5.1 Seed-based rs-fMRI analysis

rs-fMRI FC analysis was conducted using the CONN toolbox (Whitfield-Gabrieli & Nieto-Castanon, 2012) with the default_MNI preprocessed pipeline (bandpass filter: 0.01-0.1 Hz; regressed confounds: white matter, CSF and estimated participant-motion parameters). The networks of interest include the visual network (VN), sensorimotor network (SMN), the default mode network (DMN), the language network (LN), the salience network (SN), the dorsal attention network (DAN), and the frontoparietal network (FPN). The seeds for each network were based on the default seed locations provided in the CONN toolbox. To define each network, given the multiple seeds involved, the correlation maps associated individual seeds, thresholded at to p < 0.0001 (FDR corrected), were combined to generate the final network definition.

#### 2.5.2 Seed-independent rs-fMRI analysis

As assessing connectivity based on the seed-based connectivity-defined network boundaries may be somewhat circular, multiple seed-independent approaches have been included. rs-fMRI preprocessing pipeline was implemented with customized procedures based on tools from FSL (Jenkinson et al., 2012), AFNI (Cox, 1996) and FreeSurfer (Fischl, 2012). The following steps were included in the preprocessing steps: (a) 3D motion correction (FSL MCFLIRT), (b) slice-timing correction (FSL slicetimer), (c) brain extraction (FSL bet2 and FreeSurfer mri_watershed), (d) rigid body coregistration of functional data to the individual T1 image (FSL FLIRT), (e) regression of the mean signals from white-matter (WM) and cerebrospinal fluid (CSF) regions (fsl_glm), (f) bandpass filtering to obtain frequency band 0.01-0.1 Hz (AFNI 3dBandpass), and (g) spatial smoothing with a 6 mm full-width half-maximum (FWHM) Gaussian kernel (FSL fslmaths). Once the BOLD signal had been preprocessed, we computed the following metrics:

1. Resting-state fluctuation amplitude (RSFA), a measure of BOLD signal fluctuation power, calculated as the standard deviation of the BOLD signal normalized by the mean of the BOLD signal. This metric reflects the BOLD signal temporal variability that underlies the connectivity measurement.
2. lFCD, calculated using the AFNI (3dLFCD) (Cox, 1996) with a threshold of 0.6, with the neighbourhood data defined to include the 6 face-touching voxels, as suggested by the previous study (Tomasi & Volkow, 2010). gFCD, calculated using the same threshold value (0.6) but within the entire gray matter (Tomasi & Volkow, 2011). The gFCD and lFCD reflect the number of voxels that each voxel is significantly correlated to (globally or locally). The regions with high gFCD and lFCD have previously been referred to as functional connectivity hubs (Shokri-Kojori et al., 2019; Tomasi & Volkow, 2010, 2011).
3. BOLD signal entropy, calculated as the sample entropy with m = 3 and r = 0.6 related to the standard deviation as proposed by the previous study (Nezafati et al., 2020). The entropy reflects the brain’s capacity for information processing and its adaptability to various cognitive and environmental demands (Z. Wang, 2021) that may not be captured by other linear properties such as gFCD and RSFA.

Since FC density-related measurements follow an exponential distribution, log-transformed lFCD and gCDF were used in subsequent statistical analyses (Shokri-Kojori et al., 2019).

### 2.6 Baseline Perfusion Estimation

First, the ASL data was skull-stripped, and motion correction was applied to label and control images separately before surround subtraction was performed to determine the signal difference time series. To ensure the brain entered a steady state, the first five volumes of ASL data were also discarded, which is similar to in the rs-fMRI pipeline. The static CBF value was quantified using the BASIL toolbox from FSL (oxford_asl) (Chappell et al., 2009). The static CBF maps were also averaged within each functional network listed in the *Functional Network Delimitation* subsection.

Static fractional CBV will be calculated from static CBF by applying linear regression between regionally-averaged baseline static CBF (as ml/100ml/min) and CBV (as a fraction) values in gray and white matter (Eq. 1), measured by previous research with ^15^O steady-state inhalation PET in 34 healthy participants (Leenders et al., 1990; Ma et al., 2020; Zhang et al., 2018).

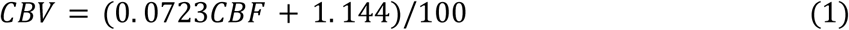

### 2.7 Baseline Metabolism Estimation

Both the SE images and the MEGRE image were first skull-stripped, and then the rigid-body coregistration was performed to align SE and MEGRE images with the T1-weighted image. Using SE images, R_2_ (refocusable transverse relaxation rate) values were determined based on a single exponential model fitted to each TE. Since the MEGRE scan is sensitive to background fields, a background field correction was performed on the MEGRE data (Kaczmarz et al., 2020). The corrected MEGRE magnitude images were then fitted using a single exponential model to estimate R *. The R ’ was calculated as the difference between R_2_ and R *. The OEF and CMRO_2_ were estimated using R_2_ ’ and perfusion parameters as described in Eq.2 and Eq.3. The theoretical details of metabolism estimation were included in our previous study (Zhong, Van Lankveld, & Chen, 2025). The static OEF and CMRO_2_ maps were also averaged within each functional network listed in the *Functional Network Delimitation* subsection.

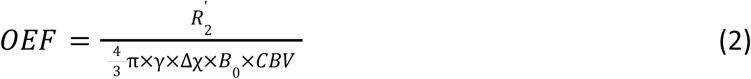

’where R_2_’ is estimated from R_2_ and R_2_*, which reflects MR signal dephasing resulting from local magnetic field inhomogeneities. γ is the proton gyromagnetic ratio, and Δχ is the susceptibility difference between oxygenated and deoxygenated blood, CBV is cerebral blood volume, and its estimation will be described in the Baseline perfusion estimation section. OEF was restricted to be between 0 and 1. Using the OEF estimate, quantitative CMRO_2_ can then be estimated (Göttler et al., 2019) as follows,

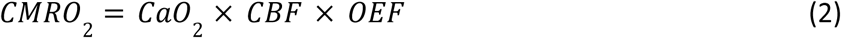

where CaO_2_ is the arterial oxygen content, set to 19 𝑚𝑙 𝑂_2_ /100 𝑚𝑙 𝑏𝑙𝑜𝑜𝑑 (An et al., 2001).

### 2.8 EEG preprocessing and analysis

#### 2.8.1 Preprocessing

EEG preprocessing was carried out using EEGLAB (version 2019.1) (Delorme & Makeig, 2004) with functions implemented in Matlab (The MathWorks Inc., Natick, Massachusetts, USA). The following steps were included in the preprocessing pipeline: 1) removal of face electrodes; 2) downsampling to 125 Hz; 3) high-pass filtering with a cut-off frequency of 1 Hz; 4) ICA denoising (removal of components related to physiological sources, including eye blinks, eye movements, residual ballistocardiograph artifacts and muscle activity); 5) channel rejection (criteria: more than 80% noisy data, line noise exceeding 4 standard deviations, flat-line segments of more than 5 seconds); 6) linear detrending.

The source reconstruction was performed using the Cartool toolbox (version 4.11) (Brunet et al., 2011). The forward model was developed for each participant using the T1-weighted MRI scans and the Locally Spherical Model with Anatomical Constraints (LSMAC) model (6-shell). The thickness and conductivity of the skull were adjusted according to each participant’s age. The inverse model was estimated using exact low-resolution brain electromagnetic tomography (eLORETA) (Imperatori et al., 2020) with 6000 solution points. The dominant orientation of the source signal was determined using singular-value decomposition (SVD) of each voxel. In addition, the sources were further mapped back to the fMRI space based on coordinates with the nearest neighbour principle so that the following metrics could be averaged with the same network ROIs as the fMRI metrics.

#### 2.8.2 EEG power

The Fourier transform was applied to the signal from each source. Band-specific fractional power was determined by dividing the power of each band by the total power across all bands, with bands defined as follows: delta (1-4 Hz), theta (4-8 Hz), alpha (8-12 Hz), beta (12-30 Hz) and low gamma (30-50 Hz). This is the same definition of bands as used in our previous study (Zhong & Chen, 2022). Unlike raw band-limited power, normalized power accounts for fluctuations in total power while quantifying the contribution of each band. Additionally, the total power across all bands was normalized by the spatial standard deviation of the total power for each participant to account for different SNRs across participants.

#### 2.8.3 EEG coherence

The imaginary part of EEG-signal coherence has been widely used to measure synchronization between two EEG signals, since it is less sensitive to volume conduction effects (Nolte et al., 2004). Our study utilized this approach to quantify EEG-based brain connectivity (Nentwich et al., 2020; Nolte et al., 2004) with a frequency resolution of 0.5 Hz and an overlap of 0.5 using scripts adapted from the Brainstorm toolbox (Tadel et al., 2011). Band-limited coherence for each band was calculated by averaging the coherence spectrum over the frequency range of each band as defined earlier. Moreover, total coherence was calculated by averaging the coherence spectrum across all bands. Here, band-limited coherence reflects functionally specific connectivity, whereas broadband coherence reflects coordination across different neuronal processes across bands.

The imaginary coherence was calculated as follows. First, we computed the cross-spectrum between the two signals, S_uv_

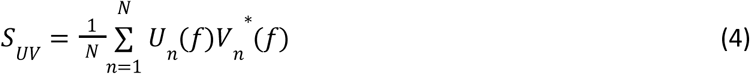

where N refers to the number of samples in the time course, and U and V refer to the Fourier transform of the two signals, respectively.

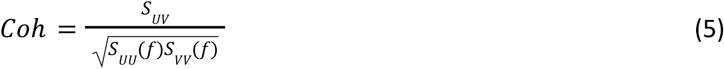

The complex coherence of the signal was calculated as the cross-spectrum normalized by their auto-spectra. Imaginary coherence is the magnitude of the imaginary part of the complex coherence.

#### 2.8.4 EEG complexity index (CI)

The complexity index was calculated as in our previous study (Zhong & Chen, 2022). The order of permutation was set to 2 and the noise threshold was 0.5 relative to the standard deviation of each band, while the scales were chosen as 3, 6, 13, 21 and 62 to represent coarse to fine temporal components of the signal. The sample entropy was used to calculate the entropy for each scale. The complexity index (CI) was calculated using the area under the multiscale entropy curve.

### 2.9 Analysis of Metabolic and Electrophysiological Underpinnings of rs-fMRI

#### 2.9.1 Analysis of associations between CMRO_2_ and EEG metrics

The associations between network-averaged metabolism and hemodynamics as well as EEG metrics were assessed using a linear mixed-effects model (LME) approach. As in our previous study (Zhong, Van Lankveld, & Chen, 2025), sex was also included as a fixed effect, since sex effects have been observed in both metabolism and hemodynamics (Aanerud et al., 2017)as well as EEG metrics (Nentwich et al., 2020; Zhong & Chen, 2022). The z-transform was applied to all parameters prior to model fitting. Outliers were detected and removed for each fitting according to the 1.5 interquartile range (IQR) criterion. An overview of the parameters of interest is presented in **Table 1** and the model (across all networks and all participants, 140 data points in total) is described as Eq. 3. A significance level of 0.05 was used with a false discovery rate (FDR).

**Table 1.** Parameters for linear mixed-effects model.

| **Y ~ 1 + X + Sex + X:Sex (Eq. 2)** |  |
| --- | --- |
| Y (EEG) | X (Metabolism and Hemodynamics) |
| Coherence (Broadband) | CMRO<sub>2</sub> |
| Coherence (Band-limited) | CBF |
| CI | OEF |
| fractional power (Band-limited) |  |
| Total power (Broadband) |  |

#### 2.9.2 Mediation analysis

Mediation analysis (**FIg. 2**) was conducted using triplets of EEG, rs-fMRI, and baseline metabolic and hemodynamic variables, considering only significant three-way associations. Prior to the analysis, the effect of sex was regressed out from all metrics. Models were implemented in MATLAB using Variational Bayesian Analysis (VBA) (Daunizeau et al., 2014) following the three-step mediation procedure of Baron and Kenny (Baron & Kenny, 1986). To assess whether the inclusion of the mediator significantly reduced the association between the independent and dependent variables (EEG to fMRI metrics), the Sobel test was applied, with significance set at p < 0.05 (Sobel, 1982). Results were classified as showing full mediation, partial mediation, or no mediation.

**Figure 2.**
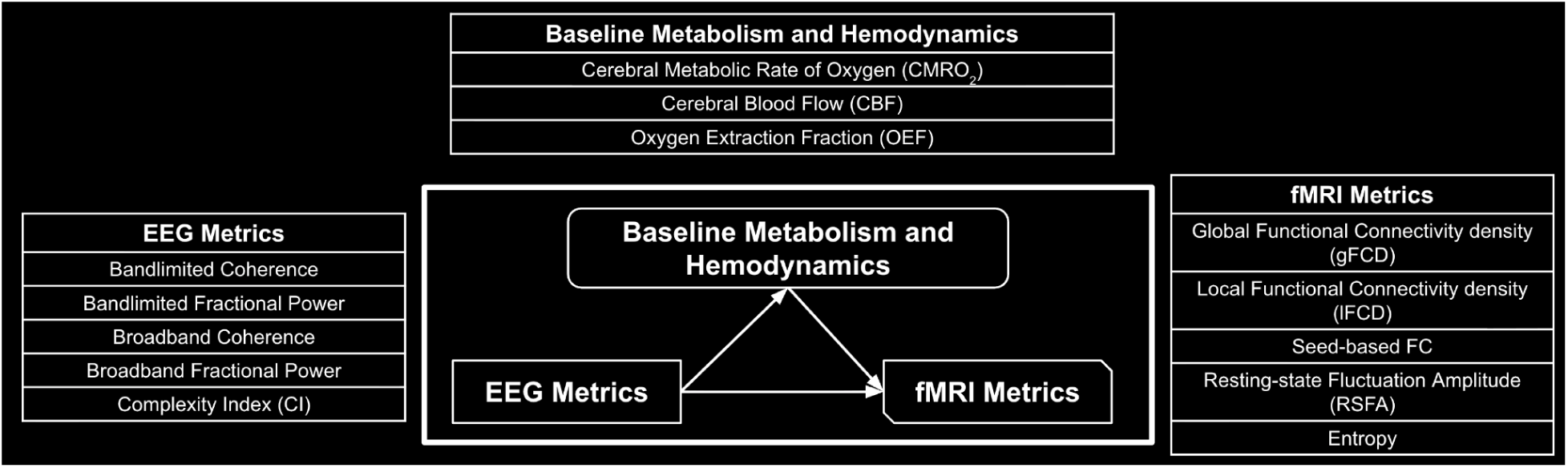
Demonstration of the meditation analysis. EEG–fMRI associations were tested for mediation by metabolic and hemodynamic metrics (CMRO_2_, CBF, or OEF).

## 3 Results

### 3.1 Metabolic and Hemodynamic Associations of Band-limited EEG Metrics

EEG band-limited fractional power was associated with baseline metabolic and hemodynamic variables, though only to a limited extent (**FIg. 3a**). Significant positive correlations were observed between CMRO_2_ and the fractional power in theta and alpha bands, while the delta band fractional power was negatively associated with OEF. Additionally, CBF showed a significant positive association with alpha band fractional power. Notably, none of the metabolic or hemodynamic metrics were significantly associated with the beta or gamma band fractional power.

**Figure 3.**
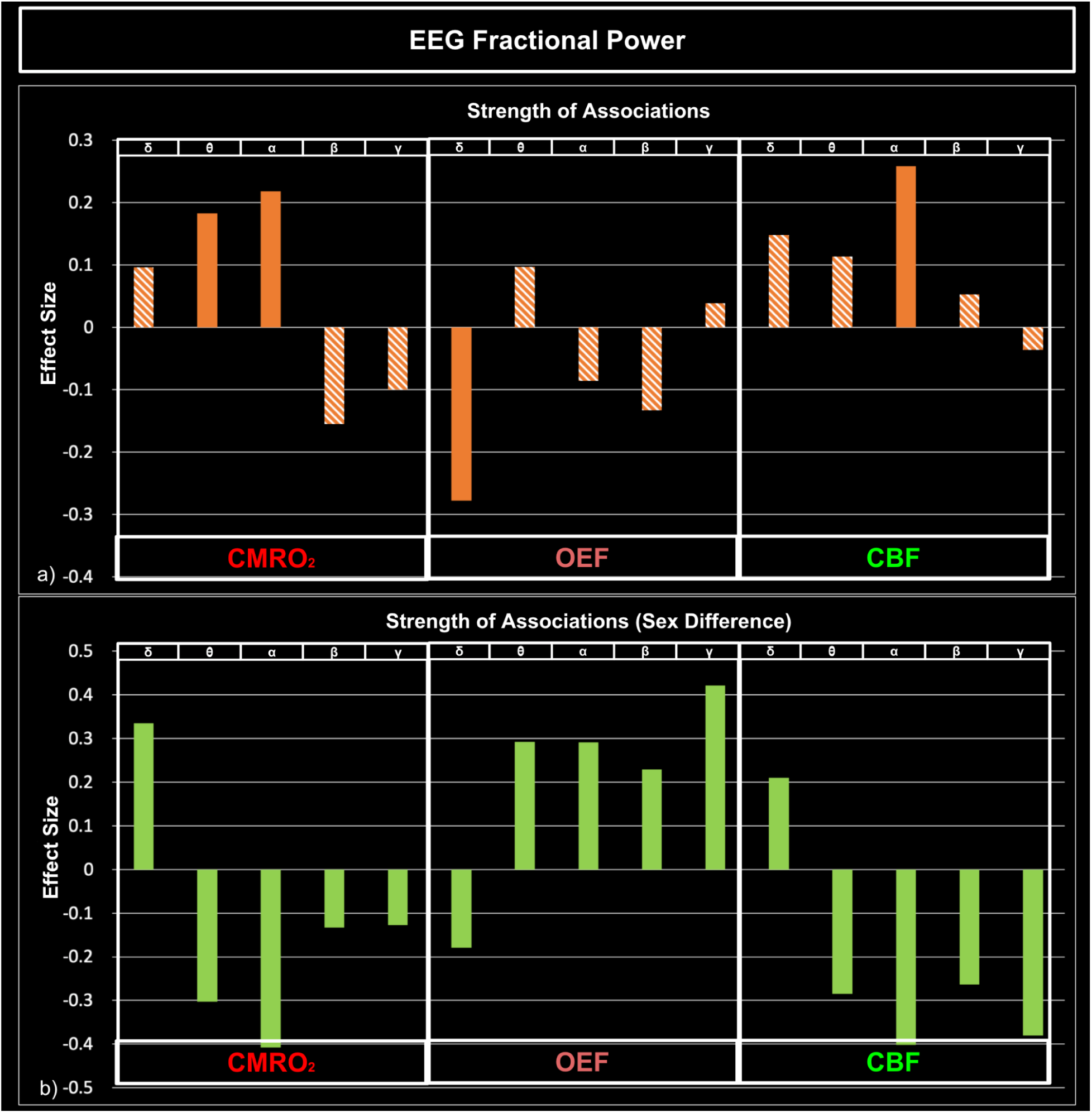
Associations between baseline metabolic and hemodynamic variables (CMRO_2_, CBF and OEF) and EEG band-limited fractional power. a) Strength of associations between rs-fMRI metrics and EEG band-limited fractional power; b) comparison of the strength of associations between males and females, with a positive effect size indicating a stronger association in males. Solid bars: association with statistical significance; striped bars: association with no statistical significance.

Significant sex interaction effects were also observed between EEG fractional power and baseline metabolic and hemodynamic variables (**FIg. 3b**). For the delta band, associations with CMRO_2_ were stronger in males, whereas associations with OEF and CBF were stronger in females. In the other frequency bands, sex interactions showed opposite patterns: females exhibited stronger associations with CMRO_2_, while males showed stronger associations with OEF and CBF. Interestingly, although many frequency bands exhibited significant sex interaction effects, some of these associations were not statistically significant. The detailed statistics are included as **Table. S1**.

CMRO_2_ and CBF were extensively associated with EEG band-limited coherences, whereas few significant associations were observed for OEF (**FIg. 4a**). Specifically, CMRO_2_ and OEF showed significant negative associations with delta, theta, beta, and gamma coherences, while their association with alpha coherence was significantly positive. CBF was significantly positively associated only with delta and gamma band coherences.

**Figure 4.**
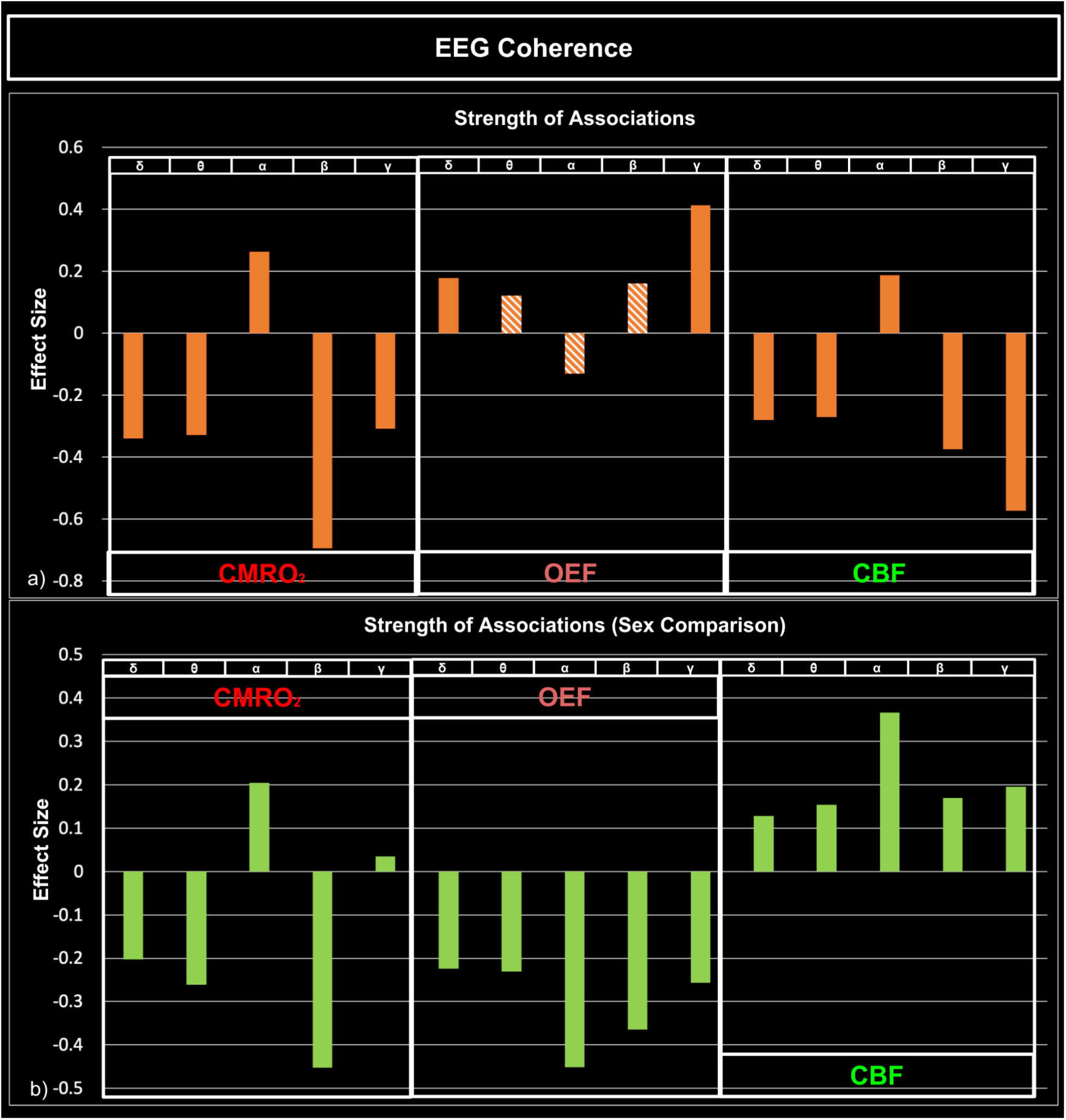
Associations between baseline metabolic and hemodynamic variables (CMRO_2_, OEF, CBF) and EEG band-limited coherence. a) Strength of associations between rs-fMRI metrics and EEG band-limited coherence; b) comparison of strength of associations between males and females, with a positive effect size indicating a stronger association in males. Solid bars: association with statistical significance; striped bars: association with no statistical significance.

Similar to band-limited fractional power, significant sex interaction effects were observed in the associations of EEG band-limited coherences (**FIg. 4b**). For CMRO_2_, females exhibited stronger associations with delta, theta, and beta coherences, whereas males showed stronger associations with alpha coherence. Females also showed stronger associations between OEF and EEG coherences across all frequency bands. In contrast, males demonstrated stronger associations between CBF and EEG coherences across all frequency bands except for the beta band. The detailed statistics are included in **Table S2**.

### 3.2 Metabolic and Hemodynamic Associations of Broadband EEG Metrics

Metabolic and hemodynamic variables were closely associated with some, but not all, EEG broadband metrics (**FIg. 5a**). Associations were primarily observed with band-limited EEG CI, but not with power and coherence. EEG CI was negatively associated with OEF and positively associated with CBF.

**Figure 5.**
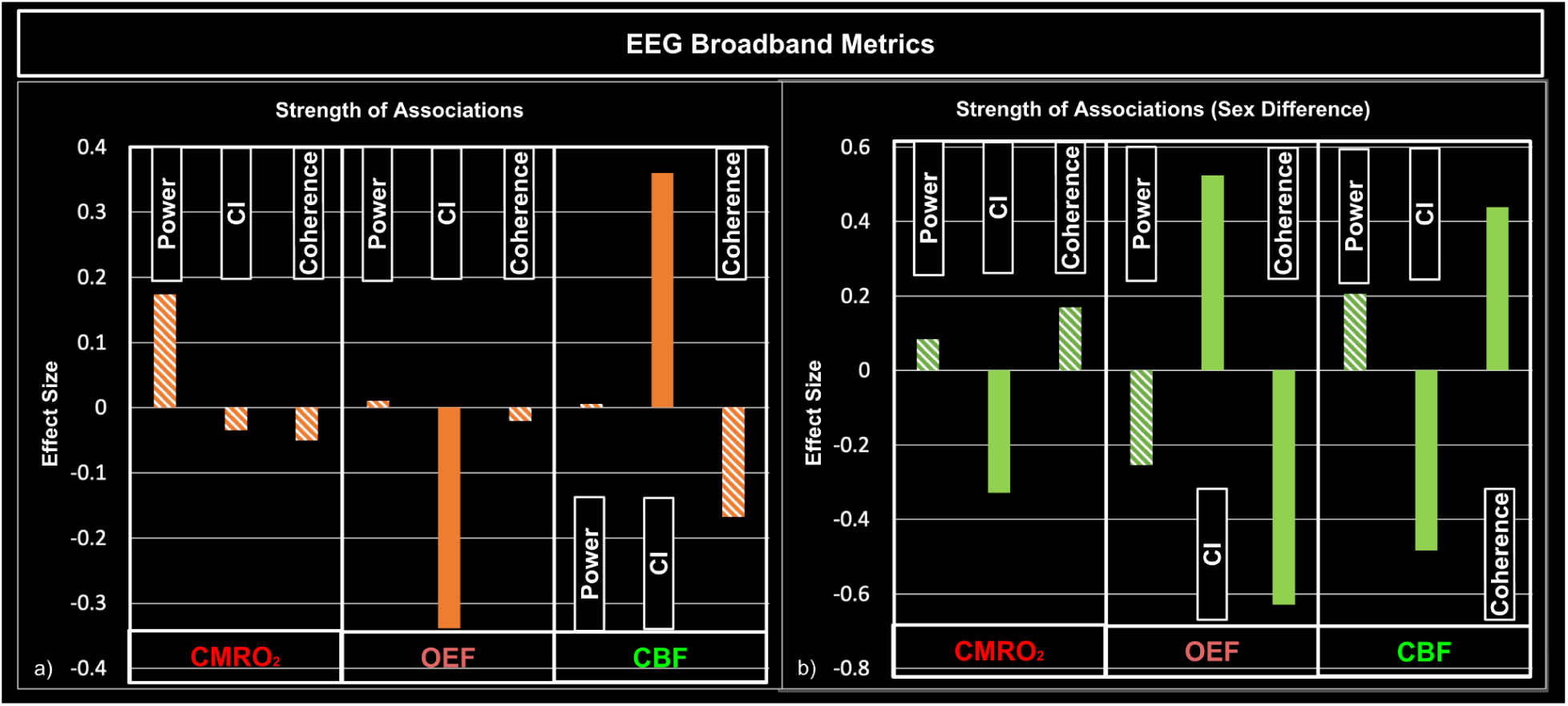
Associations between baseline metabolic and hemodynamic variables (OEF, CBF and CMRO_2_) and EEG broadband metrics. a) Strength of associations between rs-fMRI metrics and EEG band-limited broadband metrics; b) comparison of strength of associations between males and females, with a positive effect size indicating a stronger association in males. Solid bars: association with statistical significance; striped bars: association with no statistical significance.

Significant sex interaction effects were also observed (**FIg. 5b**). Females showed stronger associations with CMRO_2_ and EEG CI. CBF in females was more strongly associated with EEG CI, while in males CBF was more strongly associated with EEG coherence. Opposite to CBF, for OEF, females were more strongly associated with coherence, while males were more strongly associated with CI. The detailed statistics are included in **Table S3**.

### 3.3 The Role of Metabolism and Hemodynamics in Mediating EEG and fMRI Associations

In this work CMRO₂ was found to mediate the association between the following EEG and fMRI metrics:

● EEG power ratio: theta to lFCD (partial, ab = 0.39, p = 0.015).
● EEG coherence: delta to gFCD (full, ab = -2.66 p = 0.002); theta to gFCD (full, ab = -5.38, p = 0.009); beta to gFCD (partial, ab = -3.02, p = 0.003); beta to lFCD (full, ab = -0.58, p = 0.023); gamma to gFCD (partial, ab = -6.31, p = 0.002).

Moreover, CBF and OEF were found to mediate the association between the following EEG and fMRI metrics:

● Mediation by OEF:
  ○ EEG power ratio: none.
  ○ EEG coherence: gamma to seed-based FC (full, ab = -1.38, p = 0.011).
● Mediation by CBF:
  ○ EEG power ratio: none.
  ○ EEG coherence: beta to gFCD (partial, ab=-2.88, p=0.005); gamma to lFCD (partial, ab = 2.83, p = 0.021)

## 4. Discussion

Understanding how EEG and CMRO₂ underlie rs-fMRI metrics provides a stronger foundation for interpreting rs-fMRI findings. The main findings are:

1. Band-limited fractional power was significantly associated with baseline metabolic and hemodynamic variables.

a. Positive for CMRO_2_-theta and CMRO_2_-alpha bands;
b. Negative for OEF-delta band;
c. Positive for CBF-alpha band;
2. Band-limited coherences were significantly associated with baseline metabolic and hemodynamic variables.

a. Negative for CMRO_2_ with all bands except positive for alpha band;
b. Positive for OEF-delta and OEF-gamma
c. Negative for CBF with all bands except positive for alpha band
3. EEG broadband complex index (CI) was negatively associated with OEF, while coherence was positively associated with CBF.
4. Significant sex dependence was found for all of the above associations.
5. The baseline metabolic and hemodynamic associations with EEG significantly modulated some of the EEG-fMRI associations, primarily manifesting through CMRO_2_ but minimally through CBF.

### 4.1 The association between EEG and CMRO_2_

The association between EEG and CMRO_2_ has long been assumed (Attwell & Laughlin, 2001), yet direct evidence or quantitative assessment of this relationship has been lacking. In this study, we demonstrate how EEG fractional power and coherence (connectivity) relate to metabolic and hemodynamic measures, revealing that these associations are highly dependent on EEG oscillation frequency and differ across EEG metrics. Interestingly, nearly all networks exhibited the same polarity of association as the whole-brain results, suggesting that these associations are primarily driven by inter-participant variability rather than inter-regional variability. The physiological basis for why the EEG–CMRO_2_ association primarily emerges at the inter-participant level, even among healthy young adults, presents an intriguing question for future research.

#### 4.1.1 Band-limited Fractional Power

Fractional power has been used to reflect the strength of locally synchronized EEG activity, with stronger synchronization thought to require greater energy supply (Y. Wang et al., 2017). While this holds true in general, it does not apply uniformly across all EEG frequency bands. For low-frequency oscillations (theta and alpha), we observed clear positive associations between EEG fractional power (theta and alpha bands) and CMRO_2_ (**Fig. 3**), suggesting that increased local synchronization is accompanied by greater oxidative metabolism. In contrast, high-frequency power (beta and gamma) were not significantly associated with static CMRO_2_). To the best of our knowledge, only two studies (Buchan et al., 1997; Nagata et al., 1989) have examined the association between EEG band power and CMRO_2_ directly (rather than with CMRO_2_ surrogates such as CBF or CMRO_glu_), and both reported negative CMRO_2_ associations for EEG power below 8 Hz (delta and theta) and positive associations for power above 8 Hz (alpha and beta). However, both studies focused on patient populations (Alzheimer’s disease and cerebral infarction, respectively) rather than healthy participants. Thus, we have no healthy-adult literature as reference, suggesting that these associations may differ across physiological conditions, as also indicated by previous findings (Buchan et al., 1997). This further highlights the potential of such associations as biomarkers for neurological conditions such as Alzheimer’s disease (Buchan et al., 1997).

In theory, the EEG-CMRO_2_ associations can be explained by either EEG-CBF associations or EEG-OEF associations. For example, the positive alpha EEG–CMRO_2_ association was accompanied by a positive EEG–CBF association without a significant EEG–OEF association. This finding is consistent with prior assumptions and from task-based fMRI models, which link metabolic demand with CBF (Buxton et al., 1998; Zhong, Van Lankveld, Mathew, et al., 2025).

#### 4.1.2 Band-limited Coherence

We found that only alpha coherence was positively associated with CMRO_2_, consistent with our EEG–fMRI findings, where alpha was the only frequency band in which coherence was positively associated with gFCD (Zhong, Van Lankveld, Mathew, et al., 2025), which is in turn strongly associated with CMRO_2_ (Zhong, Van Lankveld, & Chen, 2025). This suggests that alpha coherence may dominate large-scale connectivity patterns, as discussed in our previous studies (Zhong, Van Lankveld, Mathew, et al., 2025). Unlike the complex EEG–OEF and EEG–CBF relationships observed for fractional power, the EEG–CMRO_2_ association for coherence was fully explained by the EEG–CBF association, suggesting that higher resting-state metabolism is associated with higher resting-state blood flow, consistent with relationships observed during task-evoked fMRI.

#### 4.1.3 Broadband EEG

Although CMRO_2_ is expected to reflect the overall rate of oxygen consumption across total neural activity rather than for band-specific activity, different EEG bands are likely to exhibit different metabolic profiles. Specifically, high-frequency EEG activity can reflect aerobic glycolysis (Theriault et al., 2023), which is independent from CMRO_2_, but broadband EEG metrics might still be predominantly driven by low-frequency activity (the 1/f effect) (Gyurkovics et al., 2021). Thus, a positive association between EEG broadband power and CMRO_2_ is expected. However, we did not observe significant association between broadband EEG power and CMRO_2_ after FDR correction (although a positive association was observable before FDR correction, p = 0.04), suggesting there might be a considerable role for aerobic glycolysis more broadly (**Fig. RC**). We also observed a negative CI–OEF association and a positive CI–CBF association, although the underlying mechanisms remain unclear given the complex, non-linear characteristics of CI.

### 4.3 Sex Effects on EEG-CMRO_2_ Associations

Interestingly, for EEG fractional power, the EEG–CMRO_2_ association also exhibits a pronounced sex dependence. Notably, females showed stronger EEG–CMRO_2_ and EEG–CBF associations (except for in the delta band), whereas males exhibited stronger EEG–OEF associations. These findings suggest sex differences in energy utilization patterns, potentially linked to higher aerobic glycolysis and CBF in females (Aanerud et al., 2017; Gur & Gur, 1990; Rettberg et al., 2014). In this scenario, increased neural activity that is measurable by EEG in females may shift energy metabolism from aerobic glycolysis toward oxidative phosphorylation. As discussed in our previous studies, oxidative phosphorylation has a stronger effect on CBF modulation (Zhong, Van Lankveld, & Chen, 2025), leading to increased CBF and CMRO_2_. Given the characteristically lower baseline OEF in females, as discussed earlier, an inter-subject association between CMRO_2_ and OEF in females may not exist to the same extent as in males. Although the delta band displayed the opposite trends as the other bands in terms of sex dependence, this may relate to its inhibitory role in neural processing (Amzica & Steriade, 1998; Fernández et al., 1995; Harmony, 2013). Further research is needed to disentangle these sex-dependent effects.

Strong sex-dependent effects were also observed in the associations between CMRO_2_ and EEG coherence. Similar to the case of EEG fractional power, females showed stronger associations across all frequency bands except the alpha band. This pattern may reflect a shift from aerobic glycolysis to oxidative metabolism, as discussed for fractional power. Interestingly, the sex dependencies of EEG–CBF and EEG–OEF associations differed from those observed for EEG fractional power and, in some cases, the directionalities were even reversed. For instance, females exhibited stronger EEG–OEF but weaker EEG–CBF associations. This divergence may reflect the fact that connectivity and power capture different dimensions of brain activity, and their respective associations with metabolism and haemodynamics are therefore also likely to differ.

Unlike in the case of the band-limited metrics, the sex effects for power–CBF/OEF and coherence–CBF/OEF associations were more modest. Specifically, EEG CI was the only metric manifesting significant associations with CBF and OEF and significant sex differences in these associations. This may reflect fundamental differences in the underlying interpretations of broadband (Brake et al., 2024; Férat et al., 2022), potentially in the lack of specificity in the interpretation of broadband EEG metrics. Nonetheless, the fact that we observed consistent sex differences in both bandlimited and broadband EEG metrics highlights the importance of incorporating sex differences in investigating the neurometabolic origins of rs-fMRI.

### 4.4 Metabolism and Hemodynamics Partially Mediate the EEG–fMRI Associations

The ultimate motivation for this work is to better understand the neuronal underpinnings of rs-fMRI. As discussed in our previous studies (Zhong, Van Lankveld, & Chen, 2025; Zhong, Van Lankveld, Mathew, et al., 2025), the current understanding is that the association between EEG and fMRI is indirect and likely mediated, at least in part, by CMRO_2_. Indeed, our results suggest that CMRO_2_ significantly mediates the EEG–fMRI association, primarily for gFCD and lFCD. Other metrics, such as RSFA and seed-based FC did not manifest strong EEG associations across the group. Moreover, limited mediation was observed for CBF and OEF. This is interesting given that CMRO₂ is mathematically related to OEF and CBF (Göttler et al., 2019; Zhong, Van Lankveld, & Chen, 2025), and suggests that CMRO_2_ confers unique variance not accounted for by CBF and OEF. Therefore, caution is warranted when using CBF as a surrogate for CMRO₂ across participants. However, these mediations by CMRO₂ will likely not fully explain the mechanism linking EEG and fMRI. The lack of significant CBF mediation further suggests that the CBF-CMRO_2_ coupling in a static resting state context is likely different than during task-related activity.

The basis of neurovascular coupling has long been debated. Increasingly, it is becoming recognized that neural activity can influence hemodynamics through multiple mechanisms independent of CMRO_2_ (e.g., nitric oxide signaling) (Drew, 2022). Additionally, the autonomic nervous system can modulate both EEG and fMRI signals to varying extents independently of CMRO_2_ (Bolt et al., 2025; Chalas et al., 2025), even outside brain regions directly responsible for autonomic activity. Moreover, recent insights into the role of aerobic glycolysis in brain metabolism highlight that EEG activity—particularly at higher frequencies—may not be directly linked to CMRO_2_(Theriault et al., 2023; Vaishnavi et al., 2010). However, because these high-frequency activities are strongly associated with fMRI signals (Lachaux et al., 2007), they suggest the presence of an EEG–fMRI association pathway that might not be fully mediated by CMRO_2_ (Tu et al., 2024; X. Wang et al., 2024).

### 4.5 Limitations

This study unavoidably cannot be free from limitations. Limitations and concerns directly related to the macrovascular correction, mainly involving limitations in TOF imaging acquisition and marco-VAN model simulation, have already been addressed in our previous paper (Zhong, Polimeni, et al., 2024; Zhong, Tong, et al., 2024; Zhong, Van Lankveld, & Chen, 2025; Zhong, Van Lankveld, Mathew, et al., 2025) and will not be repeated here.

Moreover, a limitation of this study is that data from the three modalities were not recorded simultaneously. As a result, we had to assume that the static properties of neuronal activity remained comparable between the EEG and rs-fMRI sessions for each participant. Thus, we used a naturalistic resting-state paradigm, presenting synchronized video content across sessions to help ensure similar brain states across modalities. Additionally, the study included a modest number of participants due to the challenges of multimodal data acquisition. Another limitation is that oxidative metabolism was not directly measured; rather, it was inferred from CBF and OEF measurements, making our CMRO₂ estimates inherently dependent on CBF. Nonetheless, this constraint is shared by many MRI- and positron emission tomography (PET)-based (Kudomi et al., 2013) approaches and remains difficult to circumvent. Finally, while we interpret the observed associations in physiological terms, they are derived from statistical associations and do not necessarily reflect causal or mechanistic relationships, particularly regarding the CMRO₂ mediation. This is a broader challenge faced by physiological MRI in general. Future studies with direct, simultaneous measurements across modalities in diverse physiological conditions are essential to establish the underlying biophysical basis of rs-fMRI from multiple perspectives.

## 5 Conclusion

Despite the widespread use of rs-fMRI, its biophysiological basis remains poorly understood. Building on our previous work, we examined how EEG metrics relate to baseline metabolism (CMRO_2_ and OEF) and hemodynamics (CBF), and further tested how EEG–CMRO_2_ associations, together with fMRI–CMRO_2_ associations, mediate the EEG–fMRI relationship. We established associations between EEG and CMRO₂ and found that CMRO₂ only partially mediates the EEG–fMRI relationship. We also uncovered substantial sex-related differences in these associations. This study provides new insight into how electrophysiological, metabolic, and rs-fMRI measures are interconnected, advancing our understanding of both rs-fMRI and cerebral physiology.

## Acknowledgement

The authors would like to acknowledge financial support from Canadian Institutes of Health Research and the Canada Research Chairs Program (JJC) and funding support from Ydessa Hendeles Graduate Scholarship (XZZ).

