## Supplemental Figures for "Linking Electrophysiological Metrics to Oxidative Metabolism: Implications for EEG–fMRI Association"

**Table S1. Statistics results for bandlimited fractional power**

| | b | p | Significance after FDR | p_thr (FDR) |
|---|---|---|---|---|
| CMRO2 vs Delta | 0.096418 | 0.31566 | FALSE | 0.0359 |
| CMRO2 vs Delta Sex | 0.33459 | 0.00067431 | TRUE | 0.0359 |
| CMRO2 vs Theta | 0.18311 | 0.029639 | TRUE | 0.0359 |
| CMRO2 vs Theta Sex | -0.30251 | 0.00044557 | TRUE | 0.0359 |
| CMRO2 vs Alpha | 0.21801 | 0.014308 | TRUE | 0.0359 |
| CMRO2 vs Alpha Sex | -0.40773 | 1.97E-05 | TRUE | 0.0359 |
| CMRO2 vs Beta | -0.15522 | 0.10565 | FALSE | 0.0359 |
| CMRO2 vs Beta Sex | -0.13262 | 0.17144 | FALSE | 0.0359 |
| CMRO2 vs Gamma | -0.099579 | 0.27101 | FALSE | 0.0359 |
| CMRO2 vs Gamma Sex | -0.12727 | 0.16777 | FALSE | 0.0359 |
| OEF vs Delta | -0.27797 | 4.16E-04 | TRUE | 0.0359 |
| OEF vs Delta Sex | -0.1789 | 0.021777 | TRUE | 0.0359 |
| OEF vs Theta | 0.097267 | 0.15841 | FALSE | 0.0359 |
| OEF vs Theta Sex | 0.29148 | 4.19E-05 | TRUE | 0.0359 |
| OEF vs Alpha | -0.085506 | 0.18613 | FALSE | 0.0359 |
| OEF vs Alpha Sex | 0.29048 | 3.72E-05 | TRUE | 0.0359 |
| OEF vs Beta | -0.13266 | 0.067949 | FALSE | 0.0359 |
| OEF vs Beta Sex | 0.2289 | 0.002158 | TRUE | 0.0359 |
| OEF vs Gamma | 0.038658 | 0.56996 | FALSE | 0.0359 |
| OEF vs Gamma Sex | 0.4208 | 1.47E-08 | TRUE | 0.0359 |
| CBF vs Delta | 0.14832 | 0.060503 | FALSE | 0.0359 |
| CBF vs Delta Sex | 0.20958 | 0.008631 | TRUE | 0.0359 |
| CBF vs Theta | 0.1132 | 0.088277 | FALSE | 0.0359 |
| CBF vs Theta Sex | -0.28456 | 3.23E-05 | TRUE | 0.0359 |
| CBF vs Alpha | 0.25862 | 2.35E-05 | TRUE | 0.0359 |
| CBF vs Alpha Sex | -0.40208 | 2.06E-09 | TRUE | 0.0359 |
| CBF vs Beta | 0.052555 | 0.47592 | FALSE | 0.0359 |

| | | | | |
|---|---|---|---|---|
| CBF vs Beta Sex | -0.26394 | 0.00057055 | TRUE | 0.0359 |
| CBF vs Gamma | -0.035727 | 0.60363 | FALSE | 0.0359 |
| CBF vs Gamma Sex | -0.37996 | 3.04E-07 | TRUE | 0.0359 |

**Table S2. Statistics results for bandlimited coherence**

| | b | p | Significance after FDR | p_thr (FDR) |
|---|---|---|---|---|
| CMRO2 vs Delta | -0.34013 | 4.22E-05 | TRUE | 0.0359 |
| CMRO2 vs Delta Sex | -0.20327 | 0.012762 | TRUE | 0.0359 |
| CMRO2 vs Theta | -0.32876 | 0.00020538 | TRUE | 0.0359 |
| CMRO2 vs Theta Sex | -0.26135 | 0.002812 | TRUE | 0.0359 |
| CMRO2 vs Alpha | 0.26242 | 0.027395 | TRUE | 0.0359 |
| CMRO2 vs Alpha Sex | 0.20414 | 0.070783 | FALSE | 0.0359 |
| CMRO2 vs Beta | -0.69364 | 3.92E-06 | TRUE | 0.0359 |
| CMRO2 vs Beta Sex | -0.45272 | 0.0014997 | TRUE | 0.0359 |
| CMRO2 vs Gamma | -0.3087 | 6.06E-03 | TRUE | 0.0359 |
| CMRO2 vs Gamma Sex | 0.035006 | 0.74079 | FALSE | 0.0359 |
| OEF vs Delta | 0.17814 | 0.0039752 | TRUE | 0.0359 |
| OEF vs Delta Sex | -0.22454 | 0.00034915 | TRUE | 0.0359 |
| OEF vs Theta | 0.12096 | 0.035946 | FALSE | 0.0359 |
| OEF vs Theta Sex | -0.23095 | 9.68E-05 | TRUE | 0.0359 |
| OEF vs Alpha | -0.12967 | 0.104 | FALSE | 0.0359 |
| OEF vs Alpha Sex | -0.45126 | 1.90E-08 | TRUE | 0.0359 |
| OEF vs Beta | 0.16135 | 0.14844 | FALSE | 0.0359 |
| OEF vs Beta Sex | -0.36517 | 0.00082332 | TRUE | 0.0359 |
| OEF vs Gamma | 0.41256 | 2.39E-07 | TRUE | 0.0359 |
| OEF vs Gamma Sex | -0.25679 | 0.00047176 | TRUE | 0.0359 |
| CBF vs Delta | -0.28001 | 7.82E-06 | TRUE | 0.0359 |
| CBF vs Delta Sex | 0.12777 | 0.035896 | TRUE | 0.0359 |
| CBF vs Theta | -0.27004 | 1.87E-06 | TRUE | 0.0359 |

| | | | | |
|---|---|---|---|---|
| CBF vs Theta Sex | 0.1536 | 0.0048802 | TRUE | 0.0359 |
| CBF vs Alpha | 0.18829 | 0.023745 | TRUE | 0.0359 |
| CBF vs Alpha Sex | 0.3659 | 6.96E-06 | TRUE | 0.0359 |
| CBF vs Beta | -0.37428 | 0.00099574 | TRUE | 0.0359 |
| CBF vs Beta Sex | 0.16926 | 0.11505 | FALSE | 0.0359 |
| CBF vs Gamma | -0.57217 | 2.97E-14 | TRUE | 0.0359 |
| CBF vs Gamma Sex | 0.19553 | 0.0023507 | TRUE | 0.0359 |

**Table S3. Statistics results for bandfree metrics**

| | b | p | Significance after FDR | p_thr (FDR) |
|---|---|---|---|---|
| CMRO2 vs Power | 0.17387 | 0.041926 | FALSE | 0.0029 |
| CMRO2 vs Power Sex | 0.083787 | 0.3258 | FALSE | 0.0029 |
| CMRO2 vs Entropy | -0.034577 | 0.70734 | FALSE | 0.0029 |
| CMRO2 vs Entropy Sex | -0.32854 | 0.00047418 | TRUE | 0.0029 |
| CMRO2 vs Coherence | -0.050164 | 0.74804 | FALSE | 0.0029 |
| CMRO2 vs Coherence Sex | 0.16964 | 2.61E-01 | FALSE | 0.0029 |
| OEF vs Power | 0.010138 | 0.87946 | FALSE | 0.0029 |
| OEF vs Power Sex | -0.25474 | 0.00020892 | TRUE | 0.0029 |
| OEF vs Entropy | -0.33829 | 7.48E-07 | TRUE | 0.0029 |
| OEF vs Entropy Sex | 0.52426 | 4.48E-13 | TRUE | 0.0029 |
| OEF vs Coherence | -0.020734 | 0.83017 | FALSE | 0.0029 |
| OEF vs Coherence Sex | -0.62854 | 3.19E-10 | TRUE | 0.0029 |
| CBF vs Power | 0.005552 | 0.9348 | FALSE | 0.0029 |
| CBF vs Power Sex | 0.20555 | 0.0029333 | FALSE | 0.0029 |
| CBF vs Entropy | 0.35959 | 2.48E-07 | TRUE | 0.0029 |
| CBF vs Entropy Sex | -0.4827 | 2.58E-11 | TRUE | 0.0029 |
| CBF vs Coherence | -0.16789 | 0.095972 | FALSE | 0.0029 |

| | | | | |
|---|---|---|---|---|
| CBF vs Coherence Sex | 0.43845 | 1.05E-05 | TRUE | 0.0029 |
